# Theory of genetic oscillations with external noisy regulation

**DOI:** 10.1101/2020.08.07.241042

**Authors:** Jose Negrete, Iván M. Lengyel, Laurel Rohde, Ravi A. Desai, Andrew C. Oates, Frank Jülicher

## Abstract

We present a general theory of noisy genetic oscillators with externally regulated production rate. The observables that characterize the genetic oscillator are discussed, and it is shown how their statistics depend on the statistics of the external regulator. We show that these observables have generic features that are observed in two different experimental systems: the expression of the circadian clock genes in fibroblasts, and in the transient and oscillatory dynamics of the segmentation clock genes observed in cells disassociated from zebrafish embryos. Our work shows that genetic oscillations with diverse biological contexts can be understood in a common framework based on delayed negative feedback system, and slow regulator dynamics.

## 1. Introduction

Embryonic cells are highly dynamic systems with time dependent biochemical states [1]. These phenotypic states are characterized by dynamical gene expression profiles. A simple but important example of dynamical gene expression is when the concentration of a protein inside a cell evolves in a cyclic and stochastic manner, this is denoted as a genetic oscillation [2]. Such dynamic process acts in many instances as a clock that determines the timing between different physiological processes [3].

Genetic oscillations can emerge from the dynamic regulation between different components of a genetic network [4]. Several theoretical studies [5, 6, 7, 8, 9] suggest that the emergence of these oscillations is mainly due to a core element that exerts time-delayed negative feedback. Such delay appears naturally inside a cell since the time it takes to produce a protein through transcription and translation is non-negligible [5, 6, 7, 10, 11]. The feasibility of this mechanism has been shown in experiments with synthetic systems [10, 11].

In this work we study a generic noisy time delayed negative feedback system *W*(*t*) externally regulated by *R*_1_(*t*) and *R*_2_(*t*) (Fig. 1). In this model *W*(*t*), *R*_1_(*t*) and *R*_2_(*t*) correspond to protein concentrations inside the cell. We develop a framework to understand the statistics that characterizes the time evolution of *W*(*t*). We find that those statistics have generic features that we observed in two different experimental cases, the circadian clock and the segmentation clock. Both systems switch stochastically between production and degradation phases with a characteristic time scale. Yet for circadian rhythms the process is sustained (Fig. 2 (a)) and for the segmentation clock the amplitude increases and decreases transiently (Fig. 2 (e)). The statistics of *W*(*t*) reflects the type of regulation that *R*_1_(*t*) and *R*_2_(*t*) exerts.

**Figure 1.**
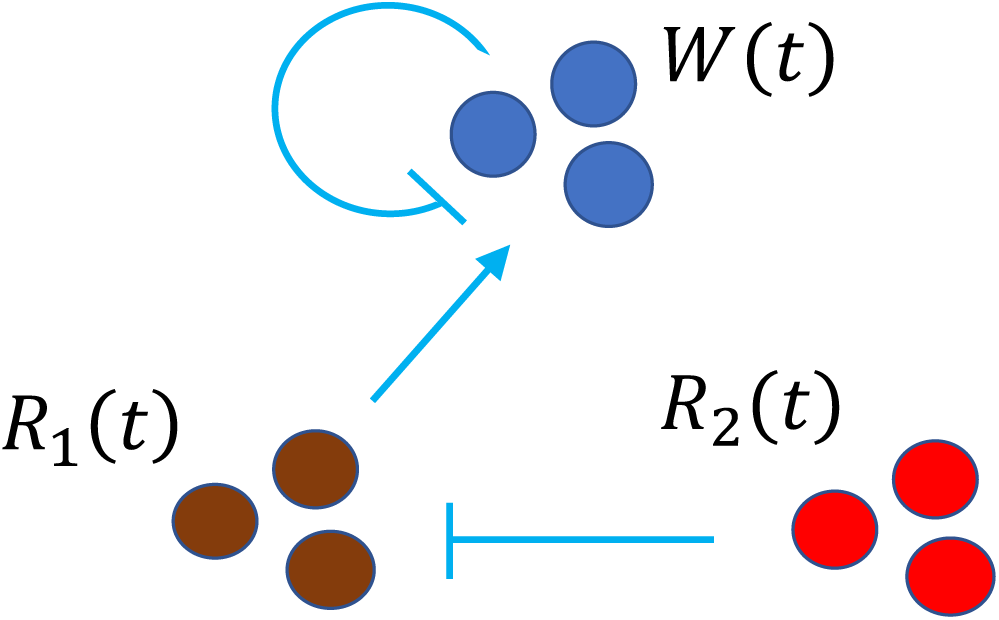
Schematic of the noisy genetic oscillator considered in this work. *W*(*t*) is the protein level of a noisy oscillating gene that is regulated by *R*_1_(*t*) and *R*_2_(*t*) which are also noisy. The model is defined by Eqns (1)-(3): *R*_1_(*t*) promotes the production rate of *W*(*t*), while a delayed negative feedback given by *W* (*t* − *T*_*d*_) represses its own production. The second regulator *R*_2_(*t*) determines when *R*_1_(*t*) is active.

**Figure 2.**
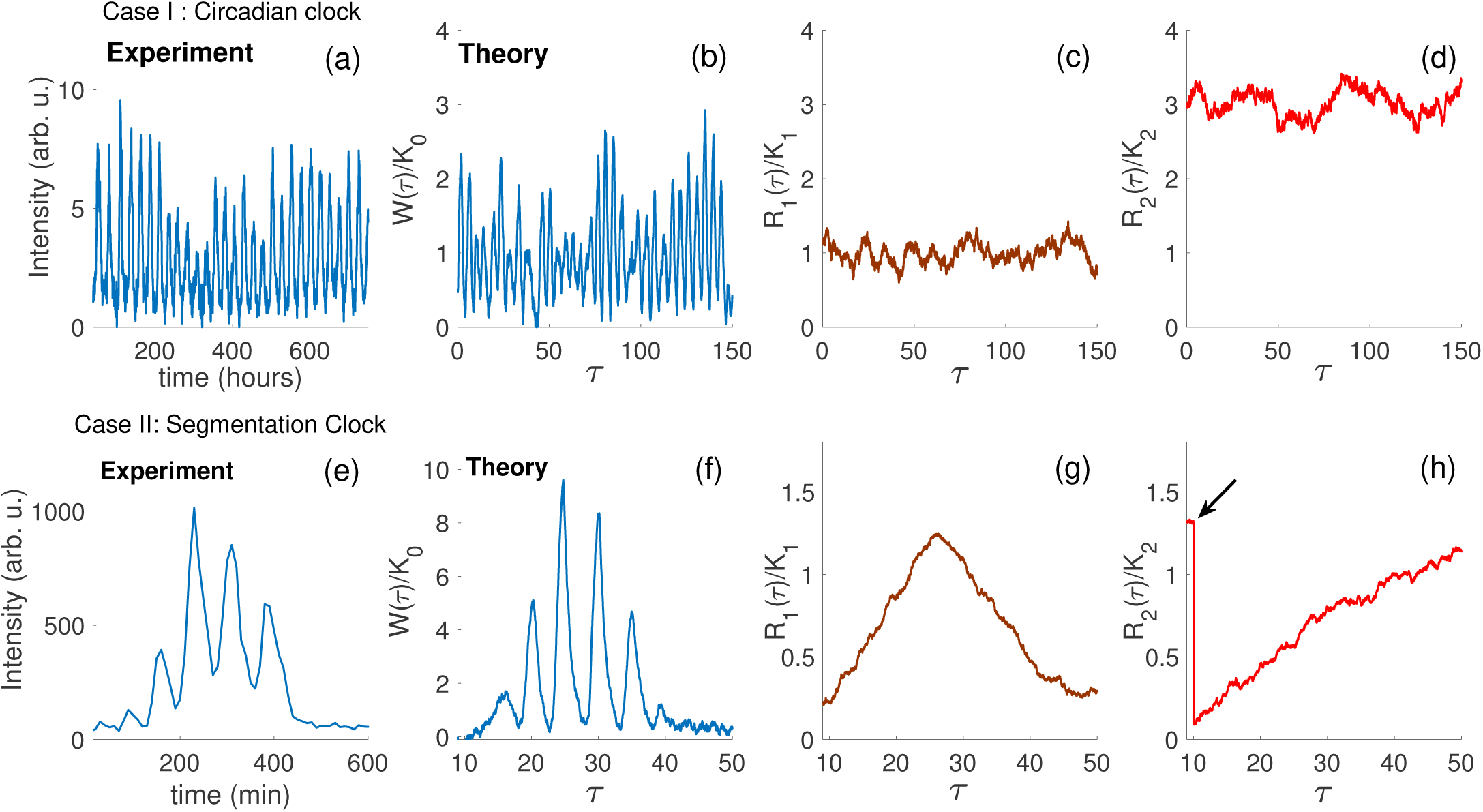
Genetic oscillations in the circadian clock and segmentation clock. Case I: Circadian clock (a-d), (a) typical trace for the time evolution in average luminescence from a fibroblast expressing PER2::luciferase (data from Leise et al. [13]). (b-d) Eqns. (1)-(3) are simulated in a regime where *W*(*t*) oscillates with a sustained amplitude regulated by *R*_1_(*t*) and *R*_2_(*t*), fluctuating around a mean value. Case II: Segmentation clock (e-h), (e) typical trace for the time evolution in average fluorescence from a cell expressing Her 1-YFP disassociated from the pre-somitic mesoderm of a zebrafish embryo. (f-h) Eqns. (1)-(3) are simulated in a regime where *W*(*t*) exhibits oscillations with an amplitude that increases and decreases transiently, *R*_1_(*t*) evolves as a pulse, which regulates the amplitude of *W*(*t*). The second regulator *R*_2_(*t*) is initially at steady state, then driven out of it to regulate the increase of *R*_1_(*t*) (time marked by a black arrow), then *R*_2_(*t*) relaxes back to equilibrium. See Table 2 for parameters.

## 2. Model of genetic oscillations with external regulation

In our model the time evolution of *W*(*t*) is given by

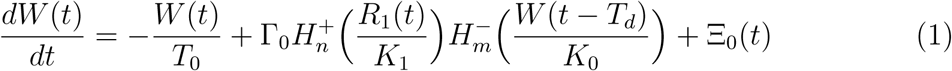

while the evolution of the regulators *R*_1_(*t*) and *R*_2_(*t*) are given by

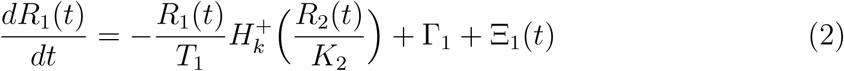

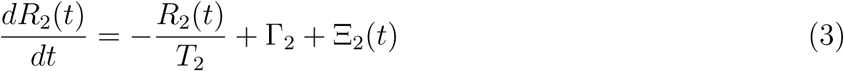

(Fig. 1). The parameter *T*_*l*_ with index *l* = {0, 1, 2} corresponds to relaxation times, Γ_*l*_ corresponds to production rates and *K*_*l*_ determines the mid-point to saturation of the Hill function 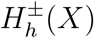, which promotes activation 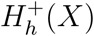 and repression 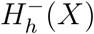 via

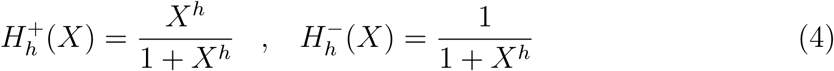

where *h* is the Hill coefficient. In a switch-like regime of production, *h* = *∞* and the Hill functions become

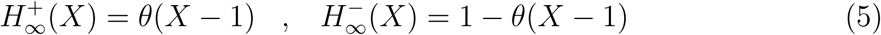

where *θ*(*X*) corresponds to the Heaviside function. The stochastic nature of gene expression in Eqns. (1)-(3) is given by the noise term Ξ_*l*_(*t*), which we have chosen to be Gaussian and correlated in time as

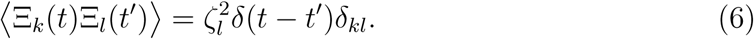

As the system evolves in time, the component *W*(*t*) switches stochastically between production and degradation phases due to gating of the production rate Γ_0_ by 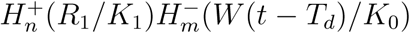 (Eqn (1)). This switching occurs with a characteristic time scale *T*_*d*_ [12] (Fig. 2 b and f).

We find that two experimental cases are captured by our model with the parameters shown in Table 2. Case I is the circadian clock, we analyzed the spatial average in luminescence intensity from single fibroblasts expressing PER2::LUC [13] (Fig. 2 a). PER2 is a component of the mammalian circadian clock which indirectly represses its own production [14]. In this case the genetic oscillator *W*(*t*) corresponds to the concentration of PER2 in a single cell (Fig. 2 b), which shows self-sustained oscillations with an amplitude that fluctuates around a constant mean value. These amplitude fluctuations are driven by the dynamics of the regulators *R*_1_(*t*) and *R*_2_(*t*), which also fluctuate around a constant mean value (Fig. 2 c-d, parameters in Table 2). Case II is the segmentation clock, which determines the sequential segmentation of the pre-somitic mesoderm into body segments in vertebrate embryos [15]. We analyzed the spatial average in fluorescence intensity from single cells expressing Her1-YFP, that were previously dissasociated from the tailbud of zebrafish embryos (Fig. 2 e, see Appendix E for experimental procedures). The protein Her1 is one of the components of the segmentation clock and also represses its own production. For the segmentation clock *W*(*t*) corresponds to the mean concentration of Her1 in a single cell, showing oscillations with an amplitude that increases and decreases transiently. In this case the oscillatory regime has specific initiation and termination times (Fig. 2 f) where the amplitude of *W*(*t*) is driven by the evolution of *R*_1_(*t*), which also increases and decreases transiently (Fig. 2 g). In turn, the evolution of *R*_1_(*t*) is determined by *R*_2_(*t*). Initially *R*_2_(*t*) is in steady state, and driven to lower values by a short pulse (arrow in Fig. 2 h). As *R*_2_(*t*) relaxes back to its steady state, *W*(*t*) and *R*_1_(*t*) return to their initial steady states.

## 3. Statistical analysis of the model

To understand the key statistics of this model, it is convenient to analyse Equation (1) in the switch-like case and rewrite it in terms of normalized concentration w = *W/K*_0_ and time *τ* = *t/T*_0_.

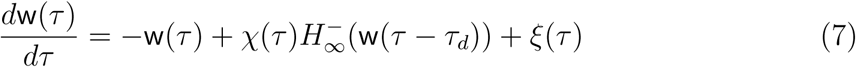

where 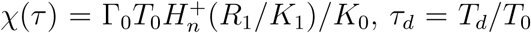 is the normalized time delay and *ξ*_*i*_(*τ*) the normalized external noise with 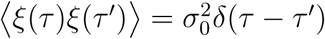 where 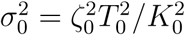. First we study the statistics of Eqn (7) when *χ* is constant. Oscillatory dynamics in w(*τ*) is observed when *χ* ≥ 1; in this regime the nonlinear term 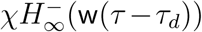 switches between two states: 0 and *χ* (Fig. 3 a). We calculate the statistics of the values of w(*τ*) at the transition points between these two states. The value at the transition 0 *→ χ* is defined as 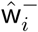, and the value at the transition *χ →* 0 is defined as 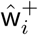 (Fig. 3 a), where the index *i* denominates the cycle number. In this case the conditional probability distribution function for the set ŵ ^*±*^ is Gaussian with

**Figure 3.**
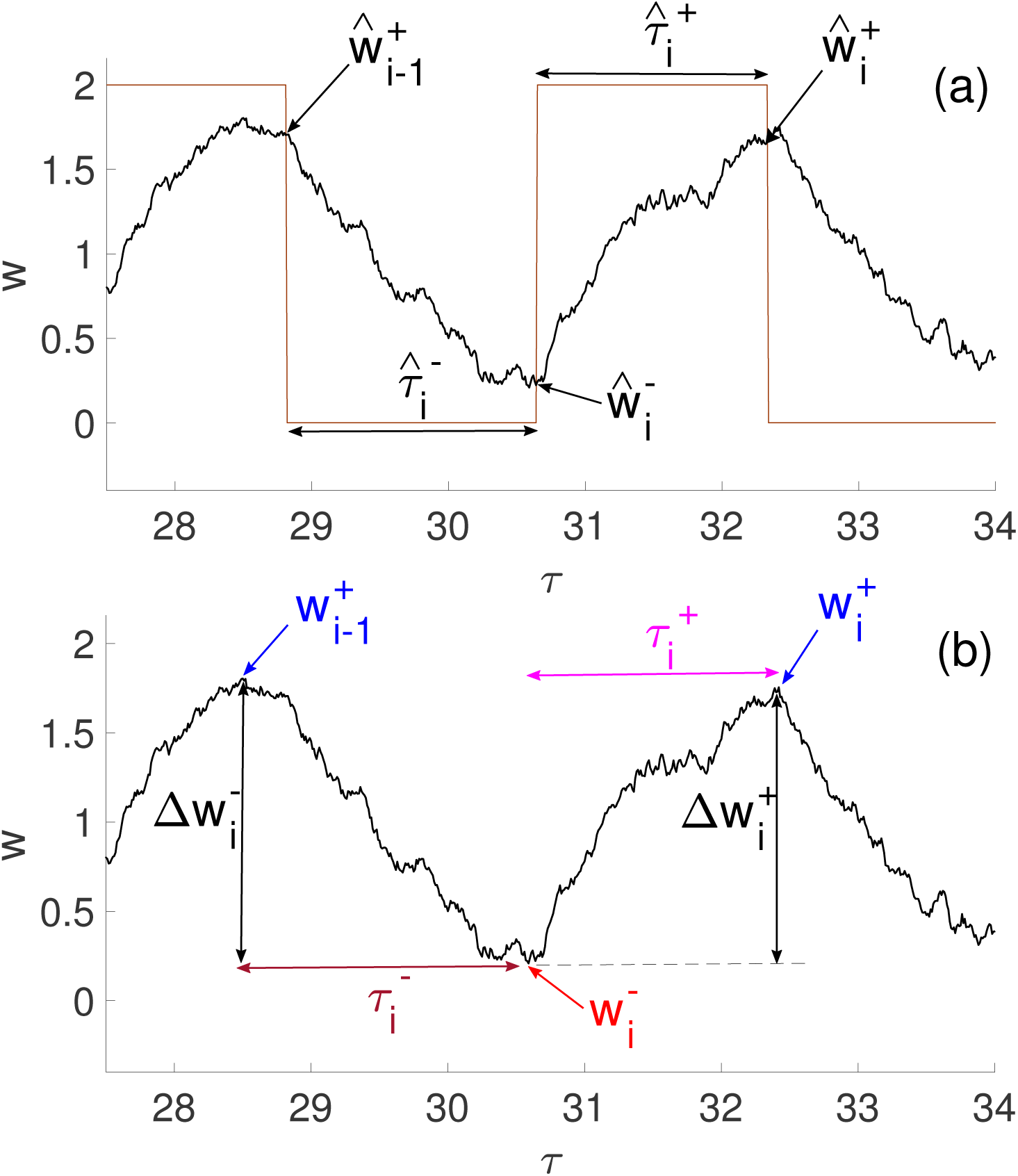
Statistical analysis of a noisy genetic oscillator. (a) In the switch-like regime of Eqn. (7) the term 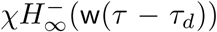 alternates between 0 and *χ* (brown line) and we determine the values of w(*τ*) at these transition points. The set ŵ ^−^ corresponds to values at the transition 0 *→ χ*, and ŵ ^+^ to values at the transition *χ →* 0. The time interval where 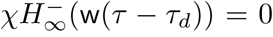 is defined as 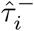 and the time interval where 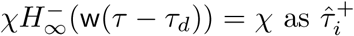. (b) In time series observed in experiment and simulation, we define proxies for ŵ ^*±*^ and 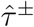 with similar statistics. We use the topographic prominence to define the set of extrema w^*±*^ that are similar to ŵ ^*±*^. The time interval between 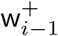 and 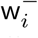 is the fall time 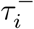, and between 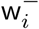 and 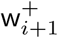 is the rise time 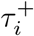. The rise and fall amplitudes 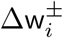 correspond to 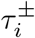.

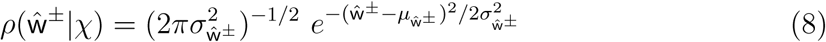

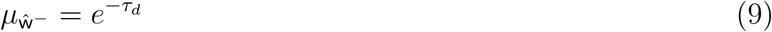

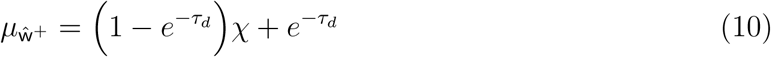

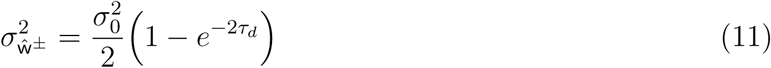

(for derivation see Appendix A). Also, we define the time intervals 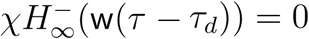 as 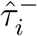 and the time intervals where 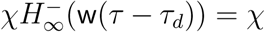 as 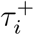(Fig. 3 a). The conditional probability distribution function for 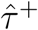 and 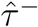 is approximately Gaussian with

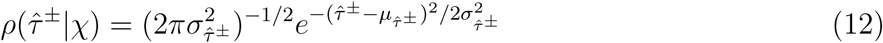

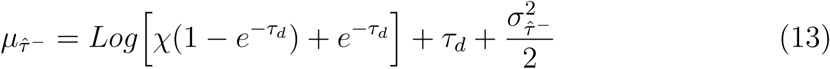

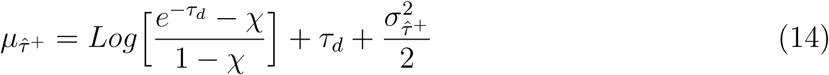

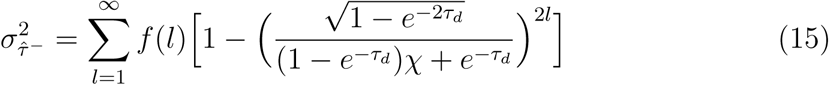

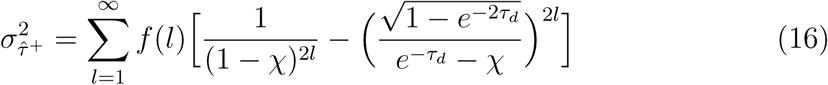

where 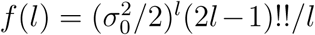(!!corresponds to the double factorial [17], see Appendix C). Note that the statistics of 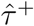 and 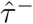 do not depend equally on *χ*, a feature that will be relevant later in the manuscript.

We can now consider the case when *χ* evolves stochastically over time. If *R*_1_(*τ*) evolves with a time scale much slower than *τ*_*d*_, then we can use the following expressions for the probability distributions for ŵ ^*±*^ and 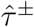

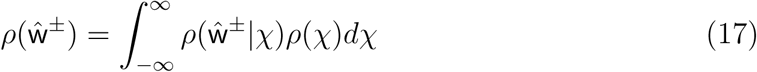

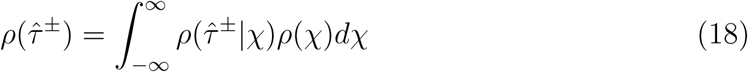

where *ρ*(*χ*) is the distribution function for *χ*. This determines the probability distributions that characterizes the noisy oscillations of our system, and now we proceed to analyze the experimental cases I and II.

## 4. Analysis of experimental cases

To analyse the experimental cases we require observables that are proxies for ŵ ^*±*^ with similar statistics and can be extracted directly from the observed time traces. We defined the set of minima w^−^ and maxima w^+^ that corresponds to the extrema at each observed oscillatory cycle (Fig. 3 b). These extrema are determined using their topographic prominence, an algorithmic measure that discerns between all local extrema and determines the most prominent peaks (see appendix B). We also define the rise 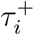 and fall 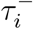 times that correspond to the time difference between points 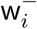 and 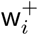, and between points 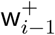 and 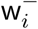 respectively. The sets *τ* ^+^ and *τ* ^−^ serve as proxies for 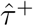 and 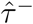. We have checked that the analytical expressions for ŵ ^*±*^ and 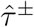 are good approximations for w^*±*^ and *τ* ^*±*^ (see appendix D). Finally, we define the rise 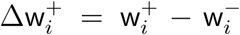 and fall 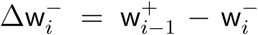 amplitudes which correspond to the time intervals 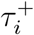 and 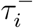 respectively.

We have been able to fit our model to both experimental cases I and II by using Equations (8)-(11) and (17). In figure 4 we plot the probability distribution functions *ρ*(w^*±*^) and *ρ*(*τ* ^*±*^) from experimental data (symbols), simulations (thick lines) and analytical approximations (dashed lines). In each case the distribution *ρ*(w^−^) is single peaked (Fig. 4 a & b), in our theory their mean values and variances are determined by *τ*_*d*_ and 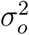 (Eqns (9 – 11)). The distribution *ρ*(w^+^) is always broader than *ρ*(w^−^) since it depends directly on the statistics of *χ* (which depends on *R*_1_(*τ*)) as shown in Eqn (10). This suggests that a Gaussian distribution for w^−^ and a broad non-Gaussian distribution for w^+^ is a characteristic of genetic oscillators. By fitting the distribution *ρ*(w^*±*^) of the model to the experimentally measured distributions, we have obtained all the parameters values in our model. Now we turn to the resulting distributions *ρ*(*τ* ^*±*^) for the rise and fall times. The distributions for *τ* ^*±*^ are similar (Fig. 4 c – d), however with distributions for *τ* ^+^ slightly skewed towards lower values, and for *τ* ^−^ skewed towards higher values. Also, the peak of *ρ*(*τ* ^+^) is at a lower value compared to *ρ*(*τ* ^−^) in each case. This suggests that the asymmetry of fast rise times and slow fall times is another characteristic of genetic oscillators.

**Figure 4.**
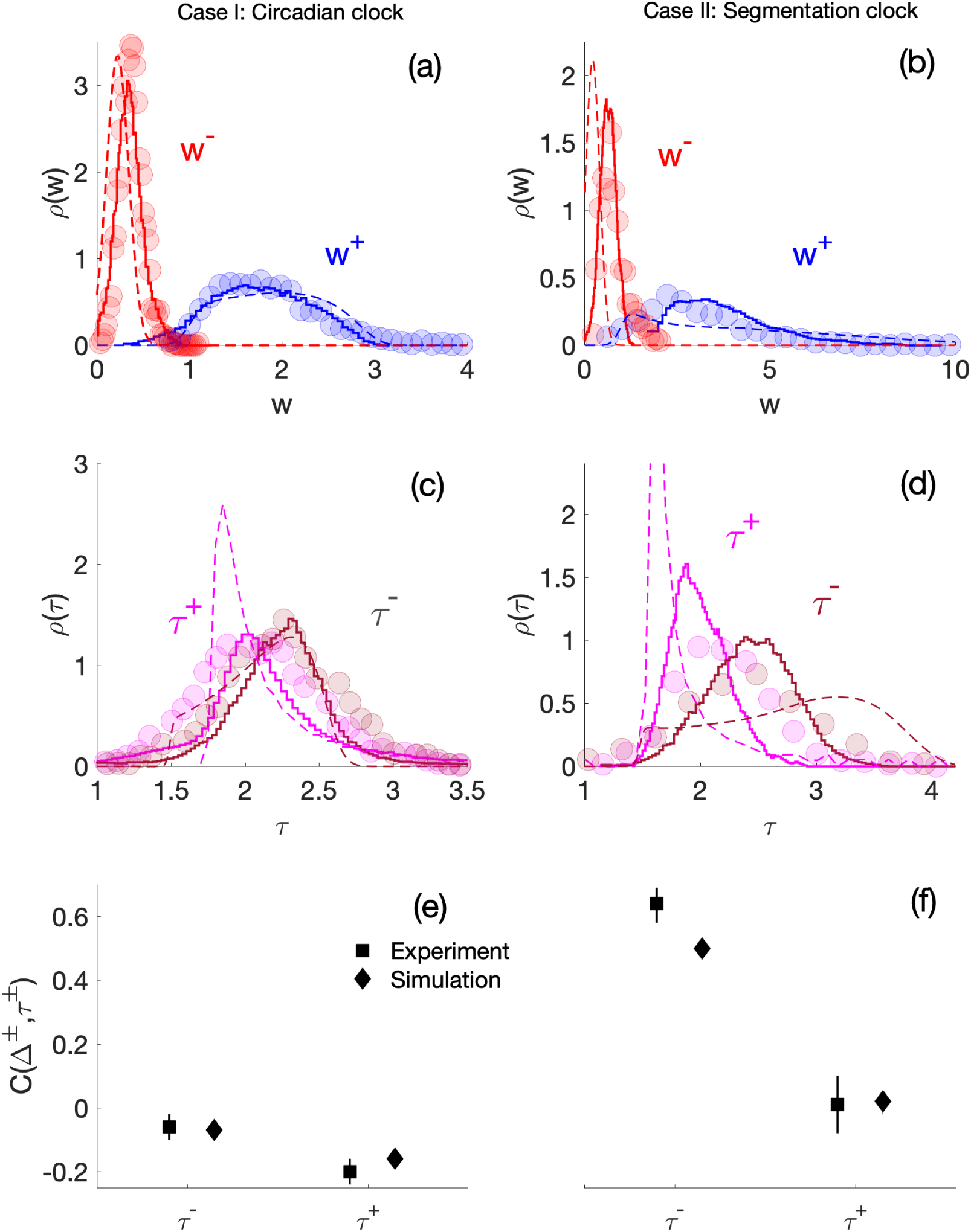
Statistics of extrema w^*±*^ and rise/fall times *τ* ^*±*^ for the circadian and segmentation clock. (a-b) Probability distributions for w^*±*^, circles correspond to experiments, thick line to simulations (parameters Table 2) and dashed lines to the analytical approximations, Eqns (8) – (11) and (17). (c-d) Distributions of the rise and fall times *τ* ^*±*^, lines and symbols as in the previous panels. The analytical approximations were obtained using Eqns (12) – (16) and (18). (e – f) The values of the linear correlation Eqn. (19) between the rise and fall amplitudes Δw^*±*^ and times *τ* ^*±*^, squares correspond to experiments and diamonds to simulations.

Finally, we note that the statistics of both ŵ ^*±*^ and 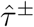 in our theory depend directly on the statistics of *χ*, therefore we looked for correlations between the rise and fall amplitudes Δw^*±*^ and times *τ* ^*±*^ in experiments and simulations. We used the following correlation coefficient to determine their correlations

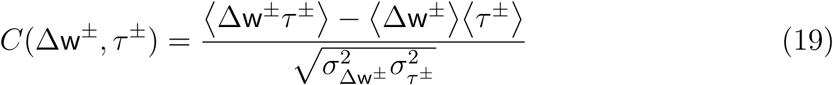

where ⟨ ⟩ denotes a steady state average. Between experiment and simulation the correlations are very similar (Fig. 4 e – f). Yet, we note that the correlations differ between Cases I and II. For Case I the correlations are weak and negative (Fig. 4 e), whereas for Case II the correlations between the fall amplitude and fall time are strong, and between the rise amplitude and rise time weak (Fig. 4 f). This difference arises because the asymmetry of fast rise times and slow fall times is amplitude dependent (Fig. 5 c). At lower amplitudes in Case I the timing is similar, whereas at higher amplitudes in Case II fall times becomes larger than rise times (Table 1).

**Table 1.** Parameter values used in simulations

| Parameter | Circadian clock | Segmentation clock |
| --- | --- | --- |
| $\Gamma_0 T_0 / K_0$ | 3.8 | 21.0 |
| $\Gamma_1 T_0 / K_0$ | 1.0 | 0.07 |
| $\Gamma_2 T_0 / K_0$ | 3.0 | 0.0667 |
| $\zeta_0^2 T_0^2 / K_0^2$ | 0.03 | 0.075 |
| $\zeta_1^2 T_0^2 / K_0^2$ | 0.0081 | 0.001 |
| $\zeta_2^2 T_0^2 / K_0^2$ | 0.0081 | 0.001 |
| $T_d / T_0$ | 1.5 | 1.5 |
| $T_1 / T_0$ | 5 | 2 |
| $T_2 / T_0$ | 5 | 20 |
| $n$ | 8 | 3 |
| $m$ | 8 | 4 |
| $k$ | 5 | 5 |

**Figure 5.**
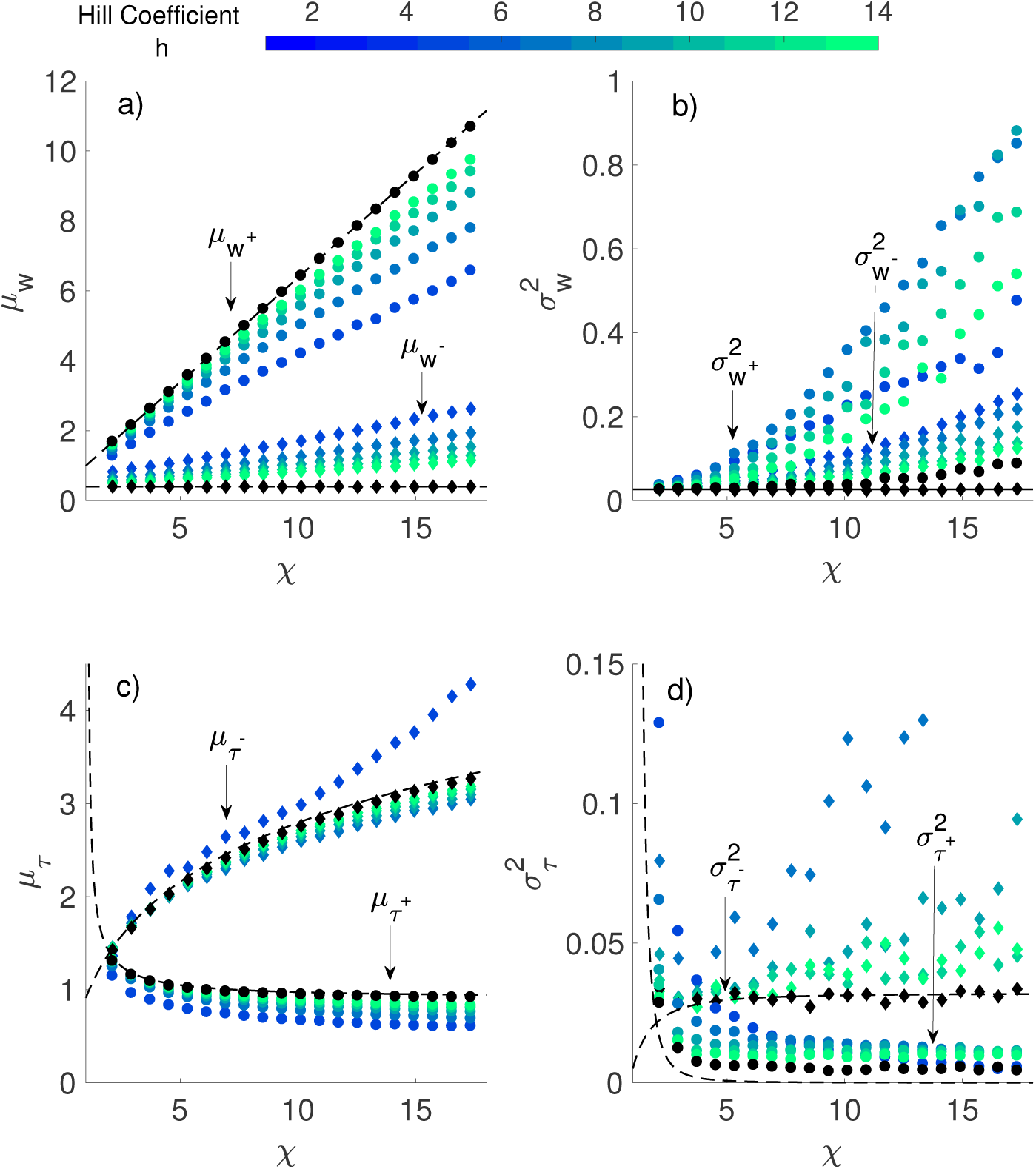
Comparison between the statistical values obtained analytically 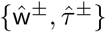 vs those from observations {w^*±*^, *τ* ^*±*^}. We simulated the steady state of Equation (1) for constant values of *R*_1_, and obtained the statistics of {w^*±*^, *τ* ^*±*^} (symbols). We also show the results from the analytical calculations for the statistics of 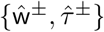 (dashed lines). (a) The mean values for w^*±*^, color circles correspond to 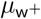 and color diamonds to 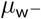. Black symbols correspond to *h* = *∞*. (b) The variance of w^*±*^, (c) the mean values of *τ* ^*±*^ and (d) the variance of *τ* ^*±*^. Color and symbols as in (a).

## 5. Conclusions

In this work we have shown that complex genetic oscillations determined by many factors [20], can be reduced to a stochastic time-delayed negative feedback oscillator with external regulation. Our model, which is motivated by Ref [12], is based on a feedback loop that switches the production of a protein alternatingly on and off. This process captures details of the oscillations of two very different genetic oscillators, the circadian clock and the segmentation clock. Both systems share similarities in the distributions of extrema of the oscillations w^*±*^, as well as the distributions of rise and fall times. The oscillations differ in the time dependence of their amplitudes, which we account for by differences in external regulation. Interestingly, the two systems differ in the correlations between amplitude and rise and fall times both in experiment and in our model. These differences reflect the different biological functions of each system. Because the circadian clock is used by the organism to measure the passage of absolute time, its period must be robust and insensitive to amplitude fluctuations [18]. This corresponds to the weak correlations between amplitude and period that we observe (Fig. 4 e). In contrast, the segmentation clock is a rhythmic patterning system, where the frequency slows down in each cell as it moves from the tailbud through the PSM while the oscillation amplitude increases [15, 19]. This corresponds to the strong correlation between amplitude and period that we observe (Fig. 4 f). This correlation arises in our model by the dependence of frequency on the amplitude, which occurs when the production rate is much larger than the degradation rate. This suggests that in the segmentation clock, the slowing of frequency occurs via the regulation of oscillation amplitude.

A key feature of genetic oscillations is that they are very noisy, with noise in both the oscillation period and amplitude. Our model can capture the statistics of these fluctuations, and can help to understand the origin of the noises in frequency and amplitude, as well their correlation. Our model highlights the importance of distinguishing between intrinsic noise, which is noise of the oscillator process itself, and extrinsic noise, which is noise in the external regulatory process.

Our analysis can provide insight in to the regulation of genetic oscillators. It is known that in mammals the circadian clock involves the production of CLOCK and BMAL1 which induce the production of CRY-PER dimers, their own repressor [14]. Our analysis reveals that amplitude fluctuations of the PER2::LUC signal are slower than the oscillation cycles. This suggests that there are pronounced but slow fluctuations associated with regulatory extrinsic noise. Because of the weak correlation between amplitude and frequency, this noise does not have a strong impact on the precision of the oscillator.

In contrast, in the segmentation clock we find that external regulation has a stronger effect on the oscillation frequency. The Her1 protein regulates its own production through negative feedback repression. The transcription factor Tbx6 is known to be an activator of Her1 production [21], and the Ripply1 protein destabilizes Tbx6 [23]. Both Tbx6 and Ripply1 are required for normal segmentation [23, 22]. Therefore in our model the activity of the regulators *R*_1_(*t*) and *R*_2_(*t*) could be analogous to concentrations of Tbx6 and Ripply1 in the embryo. Application of this theory should help to disentangle the intrinsic noise of genetic oscillators and the extrinsic noise of regulatory processes in naturally occurring systems, and help to optimise the design of synthetic systems.

## Appendix A. Statistics at turning points

In the main text the dynamics of the genetic oscillator w(*τ*) in the switch-like case are given by

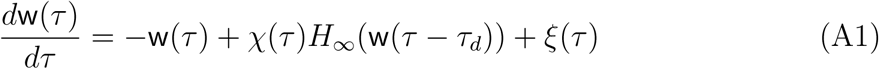

For the case when *χ* is constant we write an ansatz solution for a single realization of the form

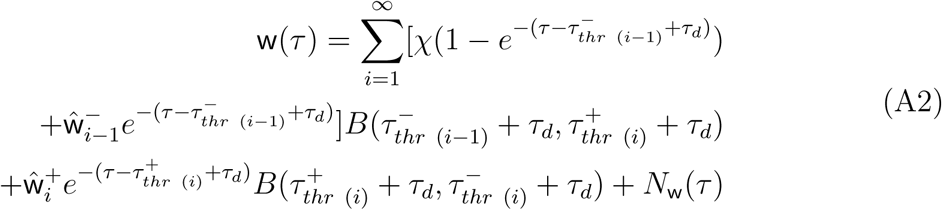

Where

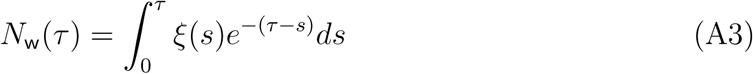

*B*(*X*_1_, *X*_2_) = *θ*(*X* − *X*_1_) − *θ*(*X* − *X*_2_), 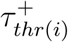 is the time when the threshold w(*τ*) = 1 is crossed upwards, and 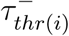 is the time when the threshold w(*τ*) = 1 is crossed downwards.

To derive the statistics of w(*τ*) at the turning points, we consider Eq. (A2) at the time points 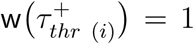 and 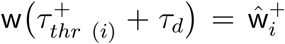. We arrive at the following expressions

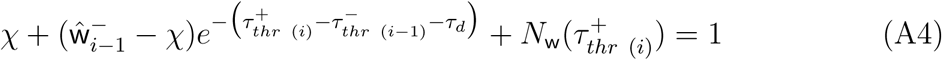

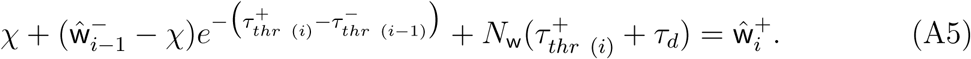

After eliminating ŵ ^−^_*i*_ we obtain

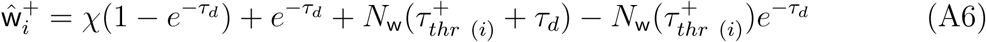

using ⟨ *N*_w_(*τ*) ⟩ = 0 the mean of ŵ ^+^_*i*_ given by Eq. (A6) is

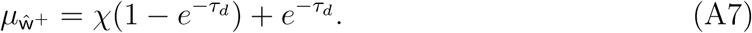

To calculate the variance we can rewrite the noise term (A6) as

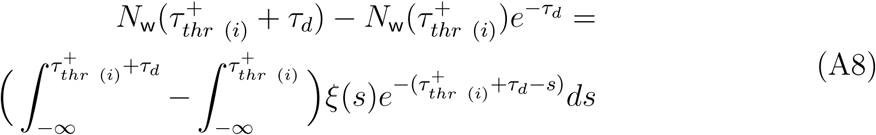

and we calculate the variance

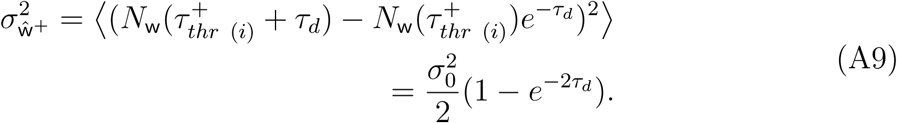

Following the same steps for ŵ ^−^ we find that the mean is given by

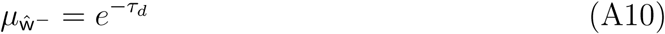

and has the same variance 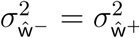.

## Appendix B. Topographic prominence

The topographic prominence *p*_*j*_ is a measure used in topography to quantify the prominence between mountain peaks. The algorithm developed by Maizlish searches the elevation of a summit relative to the highest point to which one must descend before reascending to a higher summit [16]. In this work it is used to determine which of the members of the extrema 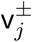 of w(*τ*) is significant enough to be part of the set 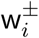. The extrema 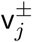 becomes part of 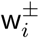 if it fulfills the condition *p*_*j*_ *> ϵ* (Fig. 6).

**Figure 6.**
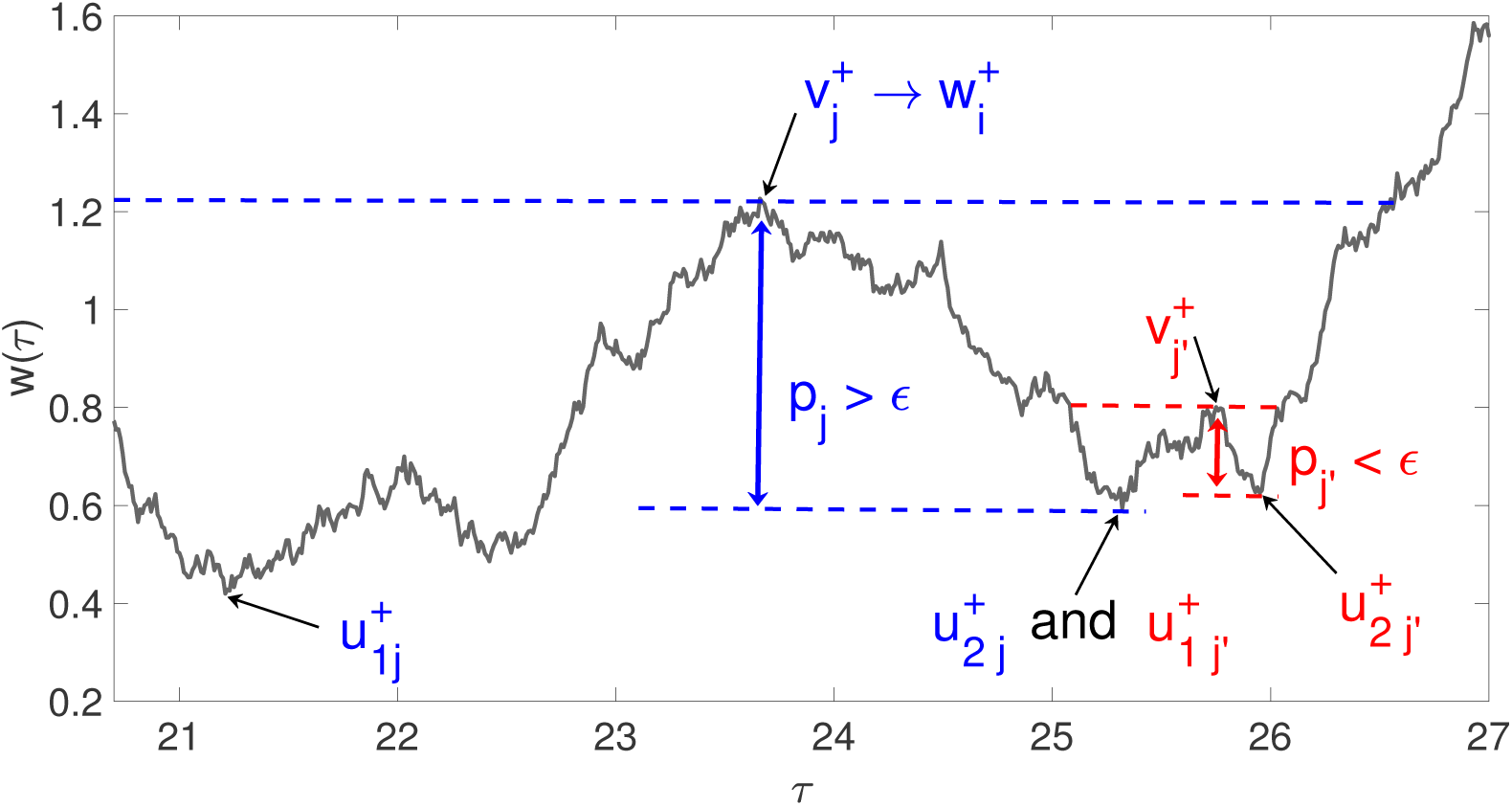
Definition of peak prominence. For each local maxima 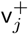, a line is shown defining a time window to the left and to the right of the extrema. We obtain the global minimum in each window and we denote 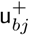 as the one with the highest value. The prominence is defined as 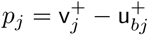.

As a first step, we assume that the signal w(*τ*) is down sampled such that we observe w(*q*Δ*t*) where the index 1 *≤ q ≤ N* and Δ*t* is the sampling time. In practice we take Δ*t* as our integration time step for numerical simulations and as the inverse sampling rate for experimental time traces. Then the algorithm to determine the topographic prominence *p*_*j*_ for each extrema 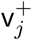 is as follows:

1. We draw a horizontal line that crosses 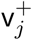, extending until it either touches the signal (red line in Fig. 6) or it reaches the end of the time series (left side of blue line in Fig. 6).
2. Two time windows are defined for the range that spans the line, one to the left of 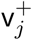 and another one to the right.
3. For each window that corresponds to each side of 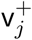, we find the global minima 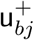where *b* = 1 corresponds to the left side and *b* = 2 to the right side.
4. We compare the values 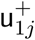 and 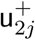 and select the highest one to calculate the topographic prominence 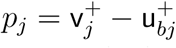.
5. Finally, we define 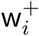 with the following rule, if *p*_*j*_ *>* ∈ then 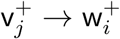 where the index *i* orders the maxima 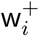 as they appear chronologically in the time series.

In Fig. 6 we have two examples showing how *p*_*j*_ is defined in each case. In these examples *p*_*j*_ and *p*_*j*_*′* share one minimum which is labeled 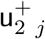 and 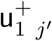 correspondingly. To define the set of 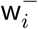 we run the same algorithm on the negative time series −w(*τ*).

## Appendix C. Statistics of timing

We start by noting two useful properties for the timing. First, the crossing times 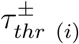 are related to the time intervals 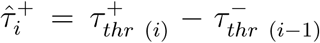 and 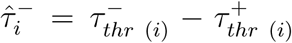. Second, we assume that 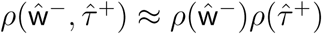 and that 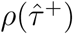 is given by a Gaussian distribution

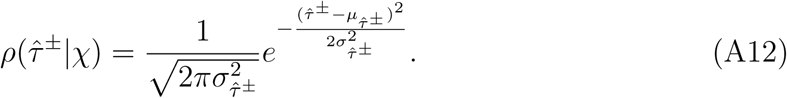

Therefore if we take the mean of Eq. (A5) we find that

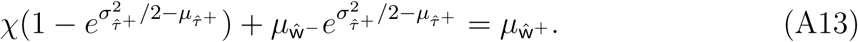

The mean and variance of 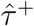 are connected by

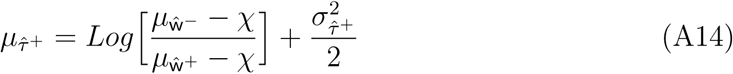

Now we can find the expression for the timing between w(*τ*) = 1 and 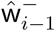 by using Eq. (A4) and rewriting 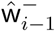 with respect to its mean and noise terms

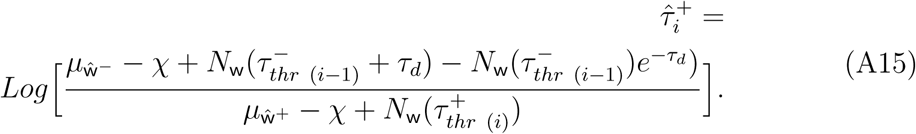

Expanding in a Taylor series for *N*_w_

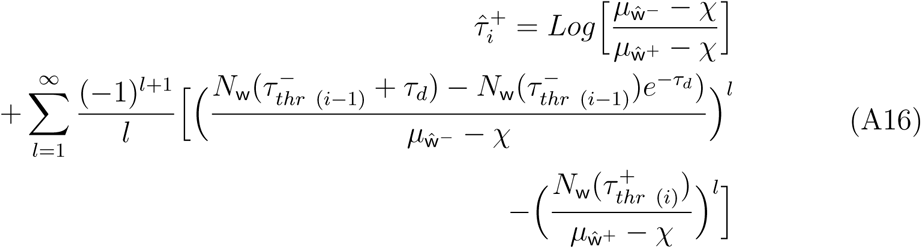

we calculate the mean and using Eq. (A14) we find that

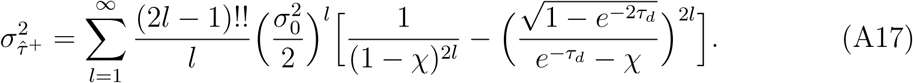

Finally, repeating the same procedure for 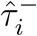 we find

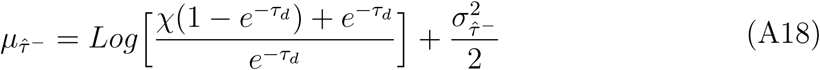

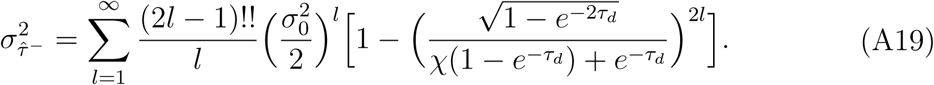

## Appendix D.

Numerical comparisons of {w^*±*^, *τ* ^*±*^} **and** 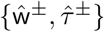

In order to compare with experiments, we need to test the proxies {w^*±*^, *τ* ^*±*^} against 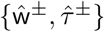. Figure 5 (a) and (b) show the comparison between the moments of *ρ*(w^*±*^) obtained from numerical calculations (black symbols) and the lines for *ρ*(ŵ^*±*^) given by Eqns. (9)-(11) for the switch like case. We note very good agreement, showing that the assumption *ρ*(w^*±*^) ≈ *ρ*(ŵ ^*±*^) is reasonable. We have calculated the statistics for the extrema for finite Hill coefficients (colored symbols). The mean value 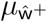 also increases monotonically with *χ*, with slope increasing as *h* increases and converges as *h → ∞*. Initially 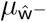 depends with *χ*, but as *h* increases its slope decreases until it reaches zero as *h → ∞*. The variances 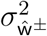 have a similar behavior. Note in Fig. 5 (c) and (d) that 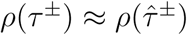 is also a good approximation.

## Appendix E. Experimental methods and signal processing

### Fibroblasts PER2::Luc (circadian clock)

We have used the data of fibroblasts with PER2::Luc label from Leise et al. [13]. The data is publicly available at https://doi.org/10.1371/journal.pone.0033334.s001, and consists in readings of the amount of photons emitted per minute from single cells. We normalized the intensity of each time trace by dividing it with its mean *I*_*norm*_ = *I/ Ī*. Since the data is filled with local minima and maxima, we have filtered each time trace with a Savitzky-Golay filter using a window of 11 time points and using a first order polynomial. The maxima and minima of each trace were found using a *peakfinder* algorithm from Matlab (MathWorks, 2016) setting a minimum peak prominence (= 0.5). We have normalized time *τ* = *t/T*_1_ in this data set using the value *T*_1_ = 5 *hrs*. The bin size of the distributions shown in figure 4 was determined using the criterium introduced by Freedman and Diaconis [24].

### Zebrafish embryos Her1-YFP (Segmentation clock)

Posterior-most paraxial mesoderm (PSM) was manually dissected from 15 somite-staged zebrafish embryos (*N* = 3 embryos and *n* = 130 cells) carrying two transgenes, Looping (Her1-YFP [25]) and Heidi (Mespbb-mkate2, unpublished). Then the tissue was dissociated into single cells, cultured without added serum or morphogens, in contrast to the work of Ref [26]. The sampling rate is set to 10 min as previously described (unpublished).

We monitored the dynamics of Her1-YFP at the single cell level. Time series of average fluorescence were extracted from each cell. Without further processing, the maxima and minima of the oscillation cycles were obtained with using the same *peakfinder* algorithm from Matlab (MathWorks, 2016). The statistics of the parameters are plotted in figure 4 (c) and (d) of the main text. We normalized time *τ* = *t/T*_1_ in this data set using the value *T*_1_ = 15 *min* and we normalized the intensity *I*_*norm*_ = *I/I*_*o*_ with *I*_*o*_ = 100 *arb*.*u*. Again, the bin size of the distributions shown in figure 4 was determined using the criterium introduced by Friedman and Diaconis.

